# Preliminary prediction of the potential distribution and consequences of *Haemaphysalis longicornis* using a simple rule-based climate envelope model

**DOI:** 10.1101/389940

**Authors:** Krisztian Magori

## Abstract

*Haemaphysalis longicornis*, the Asian longhorned (or bush) tick has been detected on a sheep in August 2017 in Hunterdon County, New Jersey. By October 26, 2018, this tick has been detected in 44 counties in 9 states along the Atlantic coast of the United States, with the first detection backdated to 2010. Here, I use a simple rule-based climate envelope model, based on a prior analysis in New Zealand, to provide a preliminary analysis of the potential range of this introduced tick species in North America. After validating this model against the counties where the tick has been already detected, I highlight the counties where this tick might cause considerable economic harm. I discuss the many limitations of this simple approach, and potential remedies for these limitations, and more sophisticated approaches. Finally, I conclude that substantial areas of the US, especially along the Gulf and Atlantic coast, are suitable for the establishment of this tick, putting millions of heads of livestock potentially at risk.

## Introduction

Invasive species cause a large amount of ecological and economic harm, due to their ability to efficiently emerge and establish in their introduced range, sometimes with devastating consequences (Pimentel et al., 2000). Specifically, invasive species that are vectors of pathogens of livestock, wildlife and humans are an even bigger concern, since they can lead to unexpected outbreaks of infectious diseases (Bonilauri et al., 2008; Akiner et al., 2016; Guichard et al., 2014). Invasive ticks are particularly difficult to control once established in an invaded area because of their small size, patchy distribution and efficient dispersal via hosts (Estrada-PeÃ±a et al., 2013).

The Asian longhorned tick (*Haemaphysalis longicornis*) is an invasive tick recently documented in North America (Rainey et al., 2018). It originates from Asia (China, Japan, Koreas, far-east Russia) and invaded the Western Pacific (Australia, New Zealand, and several Pacific Islands) (Heath et al. 2011). It is an efficient vector of several veterinary pathogens (e.g. *Theileria orientalis*, Watts et al. 2016), as well as human pathogens such as Sever Thrombocytopenia with Fever virus (STFV) (Li et al., 2016) and Rickettsia rickettsii, the causative agent of Rocky Mountain spotted fever (Lee et al., 2003). In North America, it was first detected on a sheep in Hunterdon County, New Jersey, due to extremely heavy infestation (Rainey et al., 2018). This particular tick species is able to reproduce parthenogenetically, meaning adult females can lay eggs without the presence and mating with male ticks, which can lead to extremely rapid growth and enormous densities of this tick (Hoogstral et al., 1968). The detection and identification of *Haemaphysalis longicornis* initiated increased surveillance for this tick through stakeholders such as US Fish and Wildlife Service, Rutgers University, etc. This helped identify and detect this tick at additional locations. In particular, a tick collected in 2010 off of white-tailed deer was re-examined and identified as *Haemaphysalis longicornis*, indicating that the date of introduction goes back at least to 2010 (Burtis et al., 2018). As of October 26, 2018, *Haemaphysalis longicornis* has been detected in 44 different counties, sometimes at multiple locations within a county, across 9 different states, all on the east coast of the United States (USDA-APHIS 2018).

One particular unresolved question about *Haemaphysalis longicornis* concerns its potential to geographically spread across the United States following its introduction. While all previous detections have been on the East coast, there have not been any prior studies investigating its potential distribution across the country. Agencies, such as US FWS are instructed to monitor for its presence in the entire United States, with no guideline on which part of the country it would be more likely to emerge. Knowing the potential limits for the distribution of this tick in the US would allow agencies and other stakeholders to focus monitoring and surveillance efforts in areas where this tick is most likely to establish. In addition, it would allow farmers, livestock operators and veterinarian to assess the potential for this tick to infest their livestock at particular localities, and to monitor for pathogens this tick is able to spread. Conversely, it would allow farmers and livestock owners to avoid unnecessary costs associated with monitoring and surveillance in parts of the country where it is least likely to emerge.

*Haemaphysalis longicornis*, the Asian longhorned (or bush) tick has invaded New Zealand before 1900 (Heath 2016). The tick itself caused significant economic impact due to anaemia and hide damage in livestock (Neilson 1976, Whitten 1970). *Theileria orientalis*, a tick-borne obligate intracellular protozoan parasite, causes Theileria-associated bovine anaemia (TABA), in which infected cattle can develop a severe and life-threatening anaemia (McFadden et al., 2013). The introduction of T. orientalis in 1982 and the more pathogenic Ikeda type, transmitted exclusively by *Haemaphysalis longicornis*, focused renewed attention on this tick (Vink et al., 2013). Lawrence et al. (2017) developed a simple rule-based climate envelope model in order to predict the habitat suitability across New Zealand and its potential for expansion based on future projected climate change. The rules used in this model were derived by Neilson et al. (1980) as environmental requirements based on a postal survey of sheep farmers in the Hawke’s Bay and Gisborne regions of New Zealand, correlated with local climate station data, and supported by laboratory work completed by Heath on the off-host survival of *H. longicornis* across different conditions (Heath 1974, 1977). Lawrence et al. (2017) validated this rule-based climate envelope model against a maximum entropy environmental niche model of environmental suitability for T. orientalis transmission and against a *H. longicornis* occurrence map in New Zealand.

There is still a paucity of information on the habitat requirements of *Haemaphysalis longicornis* in the US. In addition, the number of detections so far are likely to be insufficient to train a complex habitat niche model (such as MaxEnt, Phillips et al., 2006) which could discern these habitat requirements. Finally, I do not have access to fine enough scale information on the exact locations where the tick has been detected. Therefore, here I use a simplistic rule-based climate envelope model, basically adapting the work of Lawrence et al. (2017) done in New Zealand to predict the likely distribution of the Asian longhorned (bush) tick across the United States. I use publicly available climate normal data at 2.5 minute spatial resolution (Fick and Hijmans, 2017), and provide a composite habitat suitability score for *Haemaphysalis longicornis* as a surface across North America, as well as a table to summarize the average suitability of each state in the United States. I validate this prediction by overlaying the counties in which *H. longicornis* has been already detected, and provide a table of the average habitat suitability of these counties both in terms of the composite score and each of the climatic variables included. I take the mean composite habitat suitability score of the counties in which *H. longicornis* was detected as the threshold above which a county is suitable for the emergence of this tick. I then map all counties with average habitat suitability scores above this threshold that have >100,000 heads of cattle or >10,000 heads of sheep and lambs, respectively, in order to indicate counties in which the invasion of this tick is likely and could have severe economic consequences. Finally, I discuss the many limitations of this study and potential avenues towards more accurate models.

## Methods

I adapted the simple rule-based climate envelope model used by Lawrence et al. (2017) in New Zealand to predict the habitat suitability for *Haemaphysalis longicornis* across the United States. I used the statistical software R version 3.3.1 for the entire analysis (R Core Team, 2016). All code can be accessed through Github at https://github.com/kmagori/longhorned_tick. I obtained climatic variables across the United States from Worldclim Version 2 (Fick and Hijmans, 2017) at a resolution of 2.5 minute spatial resolution as raster images. The climatic variables included July minimum and July maximum monthly temperature based on 1970-2000, as well as annual mean temperature and annual total precipitation from BioClim, a part of WorldClim. Altitude was also obtained through WorldClim version 2 at the same spatial resolution. Administrative boundaries for the US, Mexico and Canada, including states and counties were obtained from GADM (Global Administrative Boundaries, gadm.org). The State of Hawaii was excluded from the analysis, which was restricted to the range of −170 ° W to −35 ° W longitude, and between 14 ° N and 90 ° N latitude. Climatic variables and altitude was cropped to this extent. I then applied a binary filter to these raster images in order to create a second set of raster images which had a value of 1 for a particular pixel if the following conditions for satisfied, one for each variable, and zero otherwise:

1. Annual mean temperature > 12 ° C
2. Minimum temperature in July >2 ° C
3. Maximum temperature in July >12 ° C
4. Annual total precipitation >1000 mm
5. Altitude <300 m

I then created a composite raster image by adding together these binary raster images to create an overall HL (*Haemaphysalis longicornis*) habitat suitability score for each pixel. Subsequently, I extracted the average HL habitat suitability score for pixels within each state of the US, along with each of the component values for those states. I identified counties where *H. longicornis* was detected and reported by October 26, 2018, and mapped it over the composite HL habitat suitability score across North America. For each of these counties, I extracted the average HL habitat suitability score as well as the average for each of its component rasters within each country. I then created a table summarizing these average scores across these counties. Finally, I averaged the HL habitat suitability score across all the counties in which *H. longicornis* was detected as a threshold value. I extracted the average HL habitat suitability score for each county within the continental US, and flagged counties with average HL habitat suitability scores above this threshold as suitable for HL.

In order to assess the potential economic impact of *H. longicornis* in the United States, I combined the above HL habitat suitability score with publicly available livestock data at the county level. I obtained the number of heads of cattle and the number of sheep and lamb per inventory for the United States at the county level in the 2012 agricultural census, using the 2012 Census of Agriculture Desktop Query Tool (USDA-NASS 2014). Data was not available for all counties of the US, in particular urban areas have not been surveyed. In addition, data in several counties were not disclosed in order to avoid reporting on individual farms. I merged the dataset containing the average HL habitat suitability scores with the datasets containing the number of cattle, as well as sheep and lambs, while making sure that county names correctly match. I then identified counties that have been marked as suitable for *H. longicornis* establishment that also had >10,000 heads of cattle, as well as >1,000 sheep and lambs, respectively, as the counties where significant economic impact from *H. longicornis* would be the most likely. Finally, I added up the total number of heads of cattle, and sheep and lambs residing in these counties suitable for *H. longicornis*, respectively.

## Results

### Distribution of HL habitat suitability score across North America

The composite HL habitat suitability score across North America is depicted on Figure 1. Areas with the highest score (5) are located primarily in the Southeastern US, south of New Jersey, and along the Gulf Coast, up to eastern Texas. To the north, the most suitable habitat for *H. longicornis* extends to central Missouri, southern Illinois and Indiana, and western Kentucky. No areas in the western US have the highest HL habitat suitability score except for a few isolated pockets in California along the coast north of San Francisco, and in the northern part of the Central Valley. The HL habitat suitability score was also the highest along both the Atlantic and the Pacific coasts of Mexico, particularly in the Yucatan peninsula, as well as on many islands of the Caribbean, particularly Cuba, the Dominican Republic and parts of Puerto Rico.

**Figure 1.**
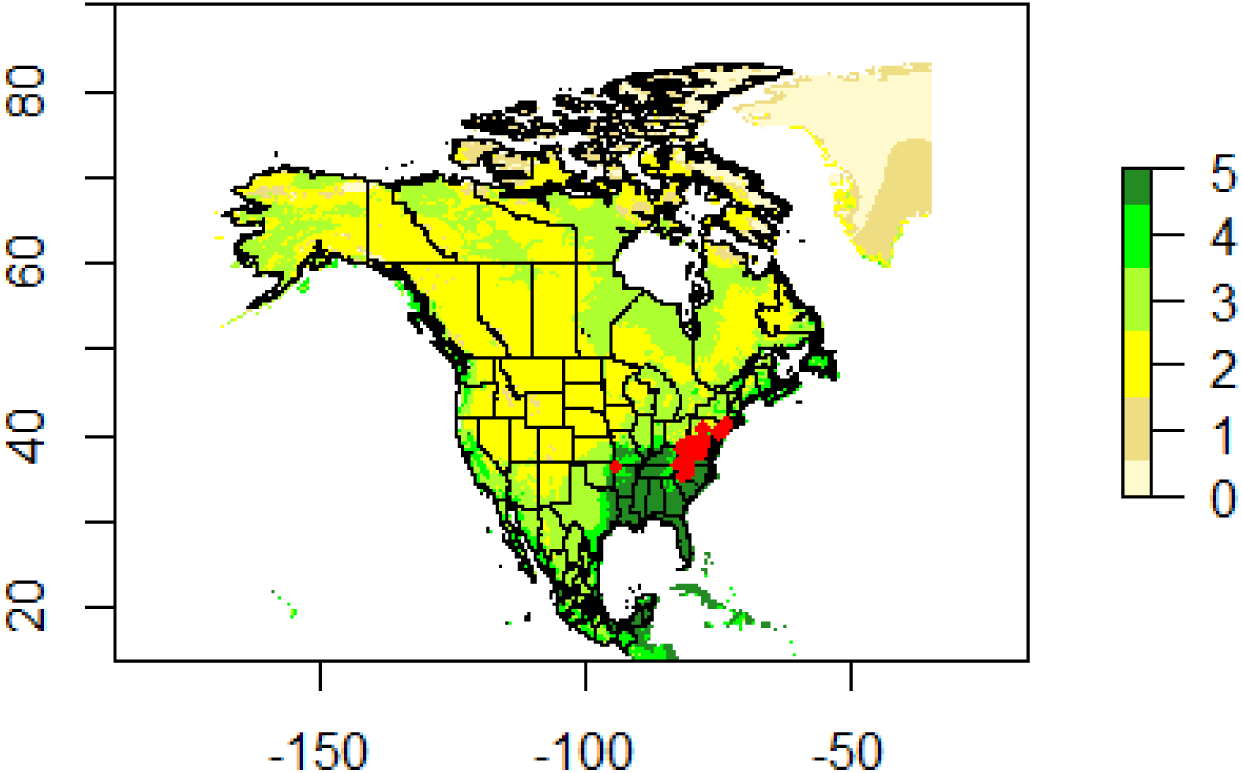
Habitat suitability of areas across North America for *Haemaphysalis longicornis*. Legend indicates colors for different levels of HL habitat suitability.

Areas with the second highest HL habitat suitability score (4) extend on the Atlantic coast of North America all the way to New Foundland and Labrador province and even isolated pixels in coastal Greenland. On the Pacific coast, this area occupies the Central Valley of California, and the coastline all the way from southern California through Oregon and Washington, as well as British Columbia, up until southern coastal Alaska, including Kodiak Island, with the addition of isolated pixels along the Columbia River between Washington and Oregon states. In Mexico, this second highest HL habitat suitability score applies to most coastal areas on both sides, as well as most areas in Central America and the Caribbean that fall into the extent.

Areas with a HL habitat suitability score of 3 include most of the Eastern US, coastal Canada, including large parts of Quebec, Ontario, Manitoba, Nunavut and Northwest Territories, as well as large parts of Alaska. On the Pacific coast of the US, the areas with an HL habitat suitability score of 3 extend somewhat more eastward than areas with an HL habitat suitability score of 4. In the southwestern US, they cover most of California, southern Nevada, large parts of Arizona, southern New Mexico, and the entirety of Texas and Oklahoma. Most of inland Mexico is in this category.

Most inland areas of the western United States fall into the category of having an HL habitat suitability score of 2. This includes eastern Washington, eastern Oregon, parts of California, most of Nevada, the entire states of Idaho, Montana, Wyoming, South Dakota, and most of Colorado, Utah, North Dakota, Nebraska, and parts of Kansas, New Mexico, Alaska and Arizona, as well as pockets in the Eastern US. In Canada, inland parts of each province fall into this category, as well as isolated areas in the mountains of Mexico.

Areas with an HL habitat suitability score of 1 occur in eastern Oregon, central Idaho, southwestern Montana and eastern Wyoming, as well as in Colorado. Areas in British Columbia and Alberta also only have an HL habitat suitability score of 1, as well as some locations in the Yukon and Northwest Territories, and in Alaska. Arctic areas of Nunavut, Greenland, and northern Alaska also have this designation. No areas in Mexico have this designation.

Finally, areas with an HL habitat suitability score of 0 only occur in isolated areas in British Columbia, Alberta, the Yukon and Northwest Territories, Nunavut, as well as Alaska. Inland areas of the arctic parts of the Northwest Territories and Nunavut, as well as Greenland have this designation. No areas in the continental US outside of Alaska, or Mexico have this designation.

### Breakdown of average HL habitat suitability score across the United States

Table 1 summarizes the average scores for HL habitat suitability for each state within the United States, along with the average values for each component, i.e. what proportion of pixels in each state satisfy each habitat requirement, organized from highest to lowest. Three of the 50 states (District of Columbia, Louisiana and Mississippi) have a perfect score of 5, meaning that every area within these states satisfy all 5 habitat requirements. Fourteen additional states (Florida, South Carolina, Delaware, Alabama, Georgia, Arkansas, North Carolina, Tennessee, Kentucky, Maryland, New Jersey, Virginia, Missouri, and Rhode Island) have average composite HL habitat suitability scores between 4 and 5. All areas within these states satisfy the July minimum and maximum habitat requirement, and portions of their areas satisfy the other habitat requirements. No part of Rhode Island satisfy the requirement that the annual mean temperature should be higher than 12, but all areas of this state satisfy all the other requirements. Another fourteen states (CT, MA, OK, IN, TX, IL, ME, NH, WV, CA, VT, NY, PA and OH) have average composite HL habitat suitability scores between 3 and 4. All areas within these states satisfy the July minimum and maximum habitat requirements (except for CA where 99.7% of the state does), and portions of their areas satisfy the other habitat requirements. No areas of CT, MA, ME and VT satisfy the requirement that the annual mean temperature should be higher than 12. Fifteen states (MI, KS, AZ, WA, WI, NM, OR, IA, AK, MN, NV, UT, ND, NE, SD) have an average composite HL habitat suitability score between 2 and 3. All areas within these states satisfy the July minimum and maximum habitat requirements (except for WA, NM, OR, and AK for July minimum, and only AK for July maximum requirement). No areas of MI, WI, IA, AK, MN, ND, NE and SD satisfy the requirement that the annual mean temperature should be higher than 12. No areas of MI, AZ, WI, NM, MN, NV, NE, UT, ND and SD satisfy the habitat requirement that total annual precipitation would exceed 1000 mm. Finally, four states (CO, MT, WY and ID) had average composite HL habitat suitability scores between 1 and 2. No areas of MT, WY and ID, and only 8% of CO satisfy the habitat requirement that annual mean temperature exceed 12. All areas within these four states satisfy the habitat requirement that July maximum temperature exceed 12, and higher than 90% of the areas of these states satisfy the habitat requirement that July minimum temperatures exceed 2. However, none of the areas of these states are lower than 300 m in altitude. Finally, none of the areas of CO, MT and WY and only 3% of ID satisfy the requirement that annual total precipitation would exceed 1000 mm.

**Table 1.**
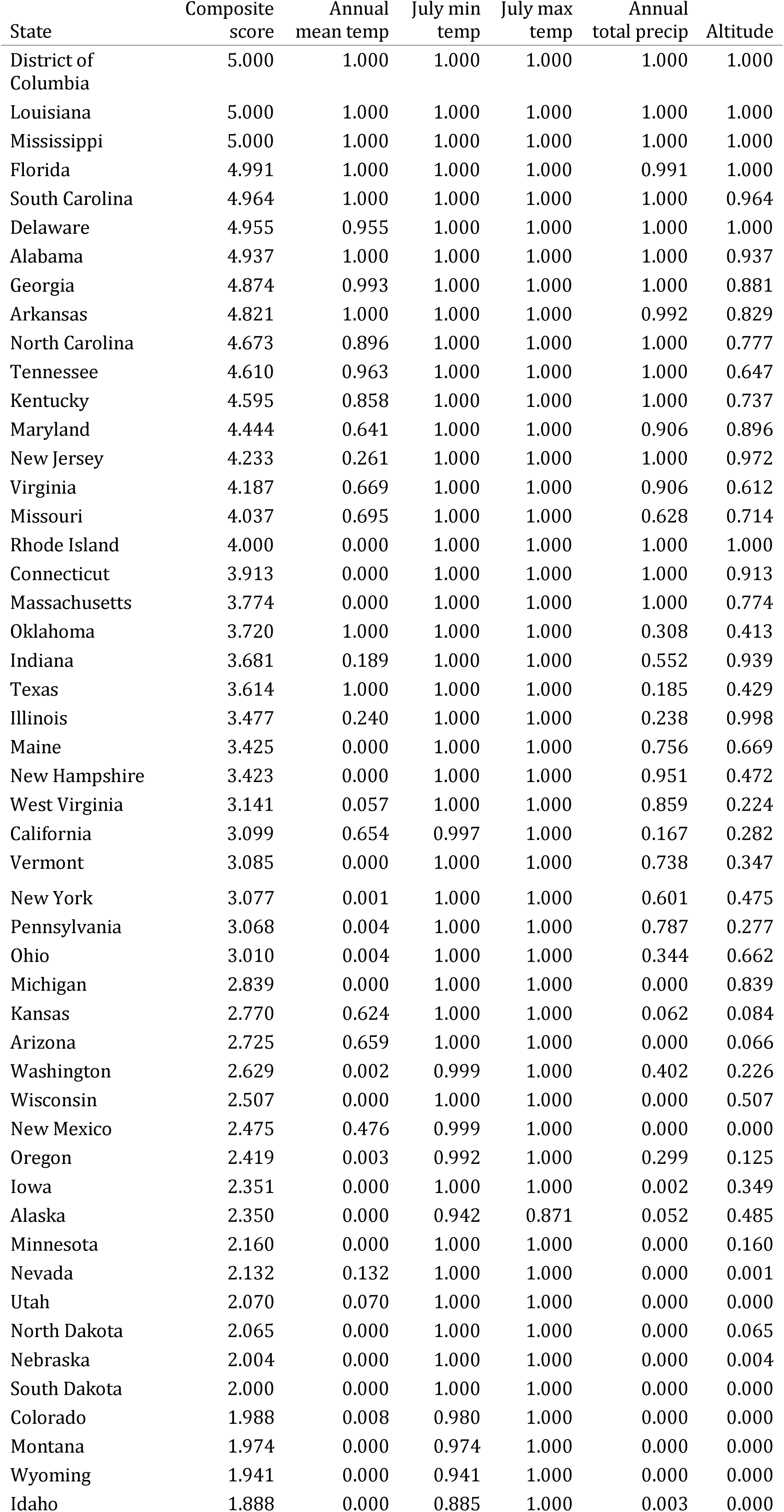
The average habitat suitability of each State of the United States for Haemaphysalis, organized from highest to lowest, as well as the scores for each component, determining the proportion of the area of each state satisfying each criteria.

| State | Composite score | Annual mean temp | July min temp | July max temp | Annual total precip | Altitude |
| --- | --- | --- | --- | --- | --- | --- |
| District of Columbia | 5.000 | 1.000 | 1.000 | 1.000 | 1.000 | 1.000 |
| Louisiana | 5.000 | 1.000 | 1.000 | 1.000 | 1.000 | 1.000 |
| Mississippi | 5.000 | 1.000 | 1.000 | 1.000 | 1.000 | 1.000 |
| Florida | 4.991 | 1.000 | 1.000 | 1.000 | 0.991 | 1.000 |
| South Carolina | 4.964 | 1.000 | 1.000 | 1.000 | 1.000 | 0.964 |
| Delaware | 4.955 | 0.955 | 1.000 | 1.000 | 1.000 | 1.000 |
| Alabama | 4.937 | 1.000 | 1.000 | 1.000 | 1.000 | 0.937 |
| Georgia | 4.874 | 0.993 | 1.000 | 1.000 | 1.000 | 0.881 |
| Arkansas | 4.821 | 1.000 | 1.000 | 1.000 | 0.992 | 0.829 |
| North Carolina | 4.673 | 0.896 | 1.000 | 1.000 | 1.000 | 0.777 |
| Tennessee | 4.610 | 0.963 | 1.000 | 1.000 | 1.000 | 0.647 |
| Kentucky | 4.595 | 0.858 | 1.000 | 1.000 | 1.000 | 0.737 |
| Maryland | 4.444 | 0.641 | 1.000 | 1.000 | 0.906 | 0.896 |
| New Jersey | 4.233 | 0.261 | 1.000 | 1.000 | 1.000 | 0.972 |
| Virginia | 4.187 | 0.669 | 1.000 | 1.000 | 0.906 | 0.612 |
| Missouri | 4.037 | 0.695 | 1.000 | 1.000 | 0.628 | 0.714 |
| Rhode Island | 4.000 | 0.000 | 1.000 | 1.000 | 1.000 | 1.000 |
| Connecticut | 3.913 | 0.000 | 1.000 | 1.000 | 1.000 | 0.913 |
| Massachusetts | 3.774 | 0.000 | 1.000 | 1.000 | 1.000 | 0.774 |
| Oklahoma | 3.720 | 1.000 | 1.000 | 1.000 | 0.308 | 0.413 |
| Indiana | 3.681 | 0.189 | 1.000 | 1.000 | 0.552 | 0.939 |
| Texas | 3.614 | 1.000 | 1.000 | 1.000 | 0.185 | 0.429 |
| Illinois | 3.477 | 0.240 | 1.000 | 1.000 | 0.238 | 0.998 |
| Maine | 3.425 | 0.000 | 1.000 | 1.000 | 0.756 | 0.669 |
| New Hampshire | 3.423 | 0.000 | 1.000 | 1.000 | 0.951 | 0.472 |
| West Virginia | 3.141 | 0.057 | 1.000 | 1.000 | 0.859 | 0.224 |
| California | 3.099 | 0.654 | 0.997 | 1.000 | 0.167 | 0.282 |
| Vermont | 3.085 | 0.000 | 1.000 | 1.000 | 0.738 | 0.347 |
| New York | 3.077 | 0.001 | 1.000 | 1.000 | 0.601 | 0.475 |
| Pennsylvania | 3.068 | 0.004 | 1.000 | 1.000 | 0.787 | 0.277 |
| Ohio | 3.010 | 0.004 | 1.000 | 1.000 | 0.344 | 0.662 |
| Michigan | 2.839 | 0.000 | 1.000 | 1.000 | 0.000 | 0.839 |
| Kansas | 2.770 | 0.624 | 1.000 | 1.000 | 0.062 | 0.084 |
| Arizona | 2.725 | 0.659 | 1.000 | 1.000 | 0.000 | 0.066 |
| Washington | 2.629 | 0.002 | 0.999 | 1.000 | 0.402 | 0.226 |
| Wisconsin | 2.507 | 0.000 | 1.000 | 1.000 | 0.000 | 0.507 |
| New Mexico | 2.475 | 0.476 | 0.999 | 1.000 | 0.000 | 0.000 |
| Oregon | 2.419 | 0.003 | 0.992 | 1.000 | 0.299 | 0.125 |
| Iowa | 2.351 | 0.000 | 1.000 | 1.000 | 0.002 | 0.349 |
| Alaska | 2.350 | 0.000 | 0.942 | 0.871 | 0.052 | 0.485 |
| Minnesota | 2.160 | 0.000 | 1.000 | 1.000 | 0.000 | 0.160 |
| Nevada | 2.132 | 0.132 | 1.000 | 1.000 | 0.000 | 0.001 |
| Utah | 2.070 | 0.070 | 1.000 | 1.000 | 0.000 | 0.000 |
| North Dakota | 2.065 | 0.000 | 1.000 | 1.000 | 0.000 | 0.065 |
| Nebraska | 2.004 | 0.000 | 1.000 | 1.000 | 0.000 | 0.004 |
| South Dakota | 2.000 | 0.000 | 1.000 | 1.000 | 0.000 | 0.000 |
| Colorado | 1.988 | 0.008 | 0.980 | 1.000 | 0.000 | 0.000 |
| Montana | 1.974 | 0.000 | 0.974 | 1.000 | 0.000 | 0.000 |
| Wyoming | 1.941 | 0.000 | 0.941 | 1.000 | 0.000 | 0.000 |
| Idaho | 1.888 | 0.000 | 0.885 | 1.000 | 0.003 | 0.000 |

### Validation of HL habitat suitability scores in counties where HL has been detected

Figure 1 depicts the counties where *H. longicornis* has been detected, overlaid on the composite HL habitat suitability score, with most of them falling outside of areas with the highest HL habitat suitability score. Table 2 summarizes the average HL habitat suitability score for each county of detection, as well as the average values for each component, organized from highest to lowest. The mean HL habitat suitability score across all counties where HL has been detected so far is 3.61, with a range from 2.21 in Pulaski County, Virginia, to 5 in Louisa County, Virginia. 20 of the 44 counties in which *H. longicornis* has been detected has an average HL habitat suitability score of 4 or higher. 15 of the 44 counties have an average HL habitat suitability score between 3 and 4, and 9of the 44 counties have an average HL habitat suitability score between 2 and 3. All areas within the 44 counties where HL has been detected satisfy the July minimum and maximum habitat requirement. No areas in 29 of the 44 counties satisfy the habitat requirement that annual mean temperature would exceed 12. All areas in 33 of the 44 counties satisfy the habitat requirement that annual total precipitation would exceed 1000 mm. Finally, all areas within 16 of the 44 counties satisfy the habitat requirement that altitude would be lower than 300 m, while none of the areas in 14 of the 44 counties satisfy this requirement.

**Table 2.**
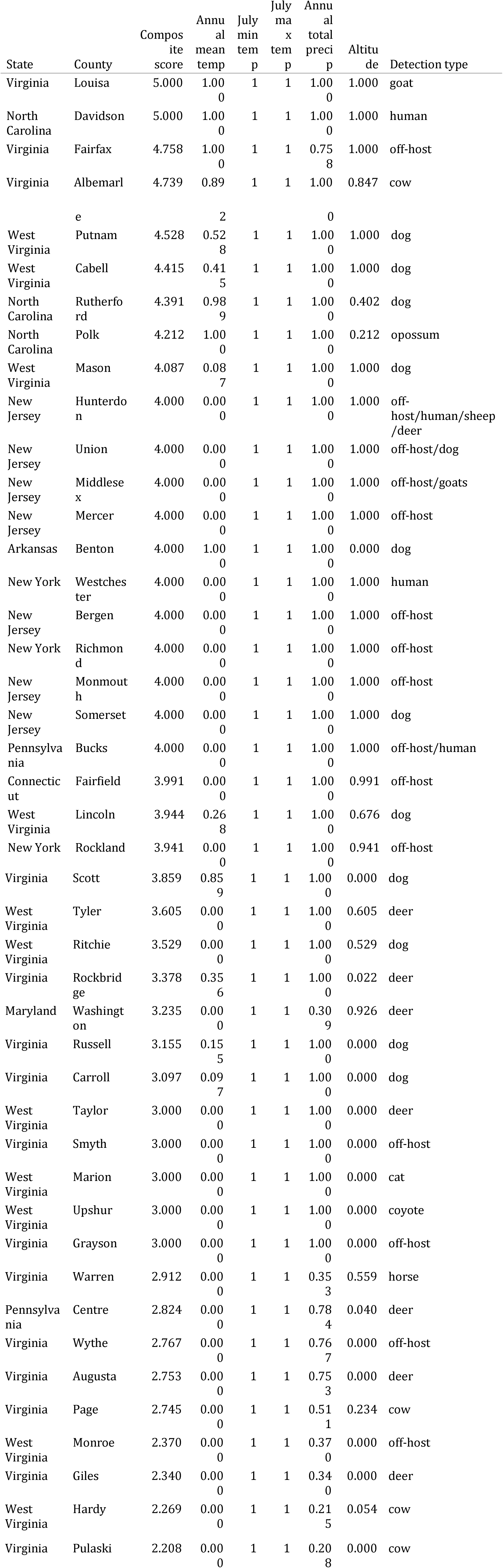
The habitat suitability score for each county where *Haemaphysalis longicornis* was detected until October 26, 2018, as well as the components, in addition to the way that the tick was collected (on or off-host, and on what host).

### Potential economic impact of HL on livestock (cattle and sheep)

Figure 2 shows the counties in which significant economic impact of *H. longicornis* is expected on cattle. These are counties that have an HL habitat suitability score above the threshold of 3.614766 (the average HL habitat suitability score of counties with documented detections) as well as at least 10,000 heads of cattle according to the 2012 USDA Census of Agriculture. There are 724 such counties across 28 states, representing 24979604 heads of cattle. Most of these counties are in the Southeastern US, from Texas to Kansas on the western edge, and all the way up to Maine of the Atlantic coast. Outside of the Southeastern US, several counties on the Pacific coast were flagged as locations where *Haemaphysalis longicornis* could establish and cause significant economic impact. In Washington State, there are 16169 heads of cattle in Clark County, WA, which has an HL habitat suitability score of 3.75; and 16631 heads of cattle in Thurston County, WA, which has an HL habitat suitability score of 3.86. In Oregon State, there are 18681, 11520 and 15365 heads of cattle in Benton, Columbia and Polk counties, respectively, which have an HL habitat suitability score of 3.73, 3.7 and 3.66. In addition, 15 counties in California (Alameda, Butte, Colusa, Contra Costa, Imperial, Kings, Marin, Merced, Sacramento, San Joaquin, Solano, Sonoma, Stanislaus, Yolo, Yuba) also have sufficient cattle numbers and are suitable for *H. longicornis*, with a total number of 2338811 cattle.

**Figure 2.**
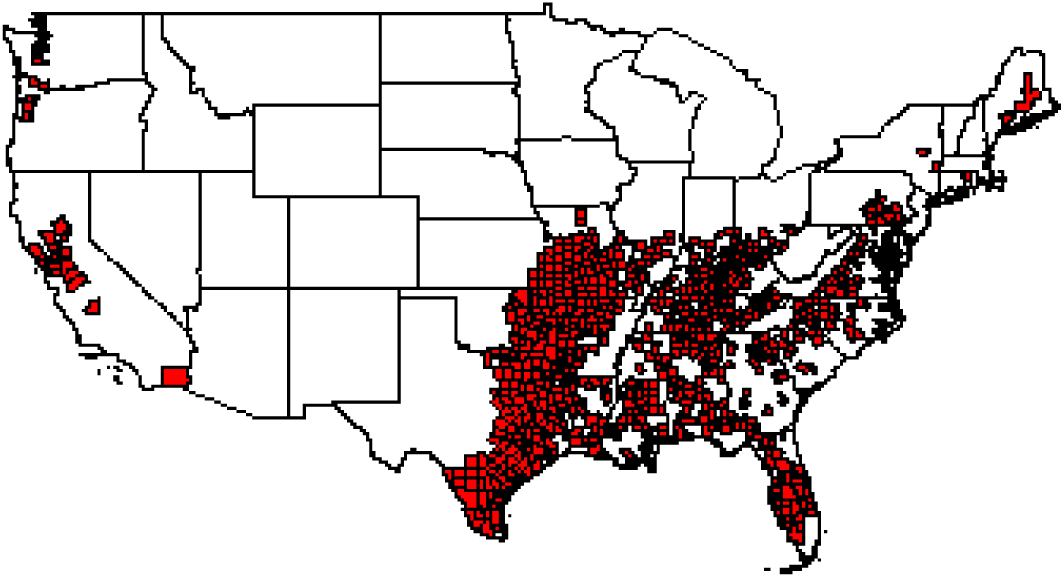
Counties in the US where substantial economic harm for cattle is likely from *Haemaphysalis longicornis*, based on the habitat suitability of these counties and the number of cattle present

Figure 3 shows the counties in which significant economic impact of *H. longicornis* is expected on sheep and lambs, where the HL habitat suitability score is above the threshold and there are at least 1,000 sheep and lambs. There are 181 such counties across 27 states, representing 564933 sheep and lamb. The general distribution of these counties is similar to the counties where economic impact is expected on cattle. Clark and Thurston counties in Washington also have 1158 and 1797 sheep and lambs, respectively. Benton, Columbia and Lincoln counties in Oregon have 3521, 1728 and 1187 sheep and lambs, respectively. The 15 counties in California also have a total number of 257603 sheep and lambs.

**Figure 3.**
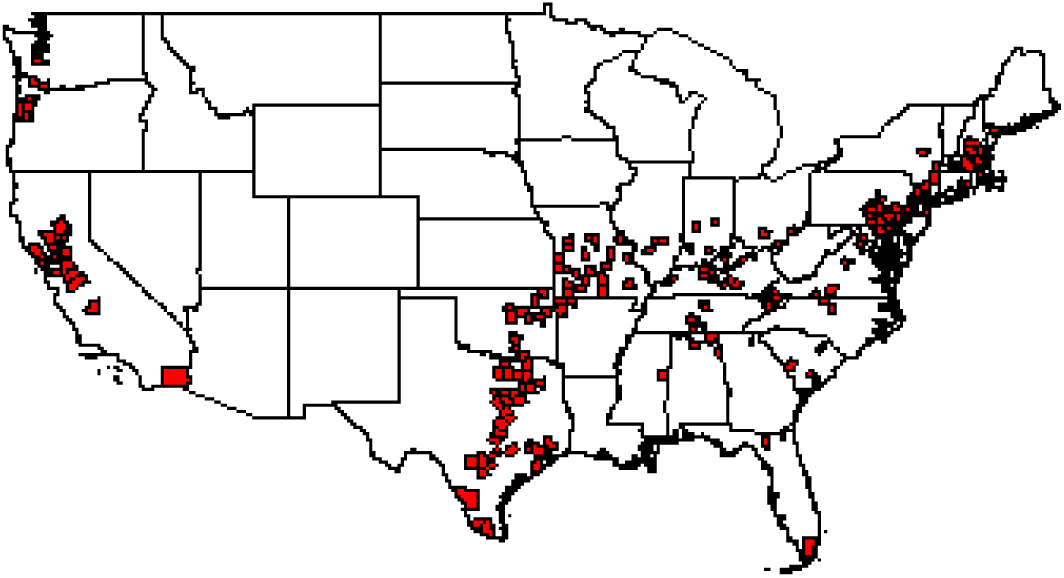
Counties in the US where substantial economic harm for sheep and lamb is likely from *Haemaphysalis longicornis*, based on the habitat suitability of these counties and the number of sheep and lamb present.

### Discussion

The above results indicate that substantial areas of North America are highly suitable (HL habitat suitability score 5) or suitable (HL habitat suitability score 4) for emergence and establishment of *Haemaphysalis longicornis*, the longhorned (bush or cattle) tick.

Specifically, in the United States, three of the 50 continental states have an average HL suitability score of 5, and 14 additional states have an average HL habitat suitability score between 4 and 5. These states are all located in the Southeastern US, the Gulf coast or the Atlantic Seaboard. Areas and states in the western US (with the exception of eastern Texas) offer generally less suitable habitat for the longhorned tick, mostly due to higher altitude, insufficient precipitation, or too cold mean annual temperature, and their combinations, with the exception of specific areas along the Pacific coast, such as the Central Valley in California. Therefore, this tick is unlikely to invade the western part of the country. However, conditions in Mexico and potentially further south in Central and South America are suitable for this tick, which could lead to its emergence and establishment there.

*Haemaphysalis longicornis* has been detected in 44 counties so far across 9 states along the Atlantic coast as well as Arkansas. The match between the habitat suitability map depicted on Figure 1 and the counties where it was detected is far from perfect. The simple rule-based climate envelope model used here was based on conditions found to describe the distribution of this tick in New Zealand, and was validated there. I am essentially extrapolating those results to North America, assuming that the conditions found to determine this tick’s distribution in New Zealand also apply across North America. There could be several problems with this approach.

First, it is possible that the strain of *Haemaphysalis longicornis* that established in North America is different from the tick that established in New Zealand. The long-horned tick has a widespread distribution across Asia, ranging from Northern China to Japan to South Korea and south to New Zealand, with different localities having different strains with different physiological requirements (Heath, 2016). Therefore, knowing the exact location where the ticks were established from would be important. In particular, if ticks from mainland Asia, such as northern China, or from Japan, were the source of the introduction, those might have very different habitat requirements from ticks in New Zealand. Second, the above analysis has been conducted at the county level. Counties are large areas that have heterogeneous distributions of climatic and other environmental variables. Ticks are sensitive and responsive to environmental variables at the micro scale (Estrada-Pena, 2003), and actively seek out suitable habitats with favorable conditions within their limited dispersal range (Estrada-Peña et al., 2013b). Therefore, the average climatic variables at the county scale might not represent the climatic variables experienced accurately-although this was not a problem in Lawrence et al. (2017), possibly due to their much higher spatial resolution (500 by 500 m). It would be possible to more accurately validate the simple rule-based climate envelope model proposed above by assessing the environmental conditions exactly where the ticks were collected within these counties-however, this information was not available to me at the time of writing.

Third, in 20 of the 31 counties where *Haemaphysalis longicornis* was detected the tick was found only on animals or people (see Table 2). The simple rule-based climate envelope model by Lawrence et al. (2017) was based on rules developed by Neilson (1980) using information on the off-host survival of *Haemaphysalis longicornis* in the lab (Heath 1974, Heath 1979). Therefore, they are not necessarily representative of conditions suitable for this tick when on host. In addition, hosts are much more mobile than the ticks themselves, some of them even crossing county boundaries. Therefore, the county where these ticks have been found on hosts might not represent the actual counties where the hosts acquired these ticks.

I elected not to adopt two of the rules used by Lawrence et al. (2017), and only applied 5 of his 7 environmental requirement in my analysis. The two rules I decided not to apply was that (1) frosts <50 per annum; and (2) Snow <=1 day per annum. There are several reasons for my choice of not using these environmental criteria. First, the second criteria was not used in the end by Lawrence et al. (2017) either, for lack of applicable data. I also did not have a readily available source of data for these environmental variables, as they are not part of the WorldClim or BioClim datasets (Fick and Hijmans, 2017). Second, frost was not well defined in Lawrence et al. (2017), so it could ity analysis that frost was not a highly important predictor of the number of farms at risk of infestation, and only included it in their model at a weight of 0.25 out of a total of 6. Third, this tick has a large distribution, including northern China and the Russian territories of Primorsky Krai, where it was documented to be able to overwinter despite cold temperatures and snow cover (Zheng et al., 2011). Prior literature indicates that while it is susceptible to freezing temperatures, this tick is able to acclimatmean either air or ground frost at various levels. Third, Lawrence et al. (2017) has shown in their sensitive to cold temperatures, increasing their glycerol and protein content (Yu et al., 2014). Finally, given that we have documented evidence that *Haemaphysalis longicornis* can overwinter in the counties where it was detected on the Atlantic coast, where we know there is frost and snow cover above these environmental criteria, it would seem counterintuitive to include these in our adopted rule-based climate envelope model.

One additional complexity is that the seasonality of this tick might be completely reversed in North America, relative to the population in New Zealand, since New Zealand is located in the southern hemisphere. Lawrence et al. (2017) used July minimum and maximum temperatures as a basis for two of their environmental rules - however, July in New Zealand is the beginning of their winter season. In that sense, one could suggest that when adopting the rule-based climate envelope model to North America, we should use December temperatures for those rules instead of July. However, very few areas in North America satisfy the requirements that December minimum temperatures would exceed 2 and that December maximum temperatures would exceed 12. Consequently, using those rules, very few areas would be considered highly suitable or suitable for the long-horned tick, including counties where it has already been detected. In addition, the origin of the ticks that were introduced to North America is at this point not known, and it is quite possible that they were introduced from areas in the Northern Hemisphere. Therefore, I did not change the month for the basis of this rule for this analysis.

Ideally, the habitat requirements of *Haemaphysalis longicornis* in the United States should be determined by analyzing the environmental conditions prevalent at the exact locations where this tick has been detected in the US, by using more sophisticated approaches such as a maximum entropy environmental niche model (Phillips et al., 2006). However, this requires fine-scale environmental data collection and knowledge of the exact location where this tick was collected off host, or where hosts acquired this tick, as well as a sufficient number of detections. The above model provides a preliminary assessment before such more sophisticated analysis could be completed, in order to inform farmers, veterinarian, agency staff involved in livestock and wildlife management, policymakers and other stakeholders. The two main findings of this analysis are that (1) there are large areas of North America, and specifically, the United States, that provide suitable conditions for *Haemaphysalis longicornis*, with significant potential risk to livestock; and (2) these suitable areas are mostly restricted to the Atlantic and Gulf coast of the eastern US, with the addition of isolated areas on the Pacific coast, such as the Central Valley of California. Predictions of the habitat suitability of specific counties, or the level of risk to livestock in those counties should be taken as preliminary results, subject to change in the future. The datasets and the code of the analysis will be updated on Github as more information becomes available and the tick is detected in additional counties. However, my hope is that these results will eventually be superseded by a much more accurate analysis based on the actual habitat requirements of this tick in its introduced range. In the meantime, I hope the reader finds this analysis informative and useful.

## References

Akiner MM, Demirci B, Babuadze G, Robert V, Schaffner F (2016) Spread of the Invasive Mosquitoes Aedes aegypti and Aedes albopictus in the Black Sea Region Increases Risk of Chikungunya, Dengue, and Zika Outbreaks in Europe. PLoS Negl Trop Dis 10(4): e0004664. https://doi.org/10.1371/journal.pntd.0004664

Bonilauri P, Bellini R, Calzolari M, et al. (2008) Chikungunya Virus in Aedes albopictus, Italy. Emerging Infectious Diseases. 14(5):852–854. doi:10.3201/eid1405.071144.

Burtis J, Egizi A, Occi J, Mader E, Lejeune M, Stafford K and L Harrington (2018) INTRUDER ALERT: LONGHORNED TICK. What you need to know about the invasive tick Haemaphysalis longicornis. https://www.acq.osd.mil/eie/afpmb/docs/bulletins/Longhorned_Tick_Fact_Sheet.pdf

Estrada-sPeña, A. (2003), The relationships between habitat topology, critical scales of connectivity and tick abundance Ixodes ricinus in a heterogeneous landscape in northern Spain. Ecography, 26: 661–671. https://onlinelibrary.wiley.com/doi/abs/10.1034/j.1600-0587.2003.03530.x

Estrada-Peña, A., & Salman, M. (2013). Current limitations in the control and spread of ticks that affect livestock: a review. Agriculture, 3(2), 221–235. http://www.mdpi.com/2077-0472/3/2/221/htm

Estrada-Peña, A., Gray, J. S., Kahl, O., Lane, R. S., & Nijhoff, A. M. (2013b). Research on the ecology of ticks and tick-borne pathogens-methodological principles and caveats. Frontiers in cellular and infection microbiology, 3, 29. https://www.frontiersin.org/articles/10.3389/fcimb.2013.00029/full

Fick, S.E. and R.J. Hijmans, 2017. Worldclim 2: New 1-km spatial resolution climate surfaces for global land areas. International Journal of Climatology. http://worldclim.org/version2

Guichard S, Guis H, Tran A, Garros C, Balenghien T, Kriticos DJ (2014) Worldwide Niche and Future Potential Distribution of Culicoides imicola, a Major Vector of Bluetongue and African Horse Sickness Viruses. PLoS ONE 9(11): e112491. https://doi.org/10.1371/journal.pone.0112491

Heath, A. C. G. (1974). An investigation into the temperature and humidity preferenda of Ixodid ticks and their distribution in relation to bioclimatic zones in Australia. Brisbane. Ph.D. thesis, University of Queensland.

Heath, A. C. G. (1979). The temperature and humidity preferences of Haemaphysalis longicornis, Ixodes holocyclus and Rhipicephalus sanguineus (Ixodidae): studies on eggs. International journal for parasitology, 9(1), 33–39. https://www.sciencedirect.com/science/article/pii/0020751979900638

Heath, A. C., Palma, R. L., Cane, R. P., & Hardwick, S. (2011). Checklist of New Zealand ticks (Acari: Ixodidae, Argasidae). Zootaxa, 2995(1), 55–63. http://www.mapress.com/j/zt/article/view/11719

Heath, A. C. G. (2016). Biology, ecology and distribution of the tick, Haemaphysalis longicornis Neumann (Acari: Ixodidae) in New Zealand. New Zealand veterinary journal, 64(1), 10–20. https://www.ingentaconnect.com/content/tandf/nzvj/2016/00000064/00000001/art00003

Hoogstraal, H., Roberts, F. H., Kohls, G. M., & Tipton, V. J. (1968). Review of Haemaphysalis (Kaiseriana) longicornis Neumann (resurrected) of Australia, New Zealand, New Caledonia, Fiji, Japan, Korea, and northeastern China and USSR, and its parthenogenetic and bisexual populations (Ixodoidea, Ixodidae). The Journal of parasitology, 1197–1213. https://www.jstor.org/stable/pdf/3276992.pdf

Lawrence, K. E., Summers, S. R., Heath, A. C. G., McFadden, A. M. J., Pulford, D. J., Tait, A. B., & Pomroy, W. E. (2017). Using a rule-based envelope model to predict the expansion of habitat suitability within New Zealand for the tick Haemaphysalis longicornis, with future projections based on two climate change scenarios. Veterinary parasitology, 243, 226–234. https://www.sciencedirect.com/science/article/pii/S030440171730300X

Lee, J., Park, H., Jung, K., Jang, W., Koh, S., Kang, S., Lee, I., Lee, W., Kim, B., Kook, Y., Park, K. and Lee, S. (2003), Identification of the Spotted Fever Group Rickettsiae Detected from Haemaphysalis longicornis in Korea. Microbiology and Immunology, 47: 301–304. doi:10.1111/j.1348-0421.2003.tb03399.x https://onlinelibrary.wiley.com/doi/abs/10.1111/j.1348-0421.2003.tb03399.x

Li Z, Bao C, Hu J, Liu W, Wang X, Zhang L, et al. (2016) Ecology of the Tick-Borne Phlebovirus Causing Severe Fever with Thrombocytopenia Syndrome in an Endemic Area of China. PLoS Negl Trop Dis 10(4): e0004574. https://doi.org/10.1371/journal.pntd.0004574

McFadden, A., Pulford, D., Lawrence, K., Frazer, J., van Andel, M., Donald, J., & Bingham, P. (2013). Epidemiology of Theileria orientalis in cattle in New Zealand. In Proceedings of the Society of Dairy Cattle Veterinarians of the NZVA Annual Conference (pp. 207–217).

Neilson, F. J. A. (1976). Cattle tick infestation in lambs. In Proceedings of the 6th Annual Seminar of the Society of Sheep and Cattle Veterinarians, New Zealand Veterinary Association (pp. 21–6).

Neilson, F. J. A. (1980). An investigation into the ecology, biology, distribution and control of Haemaphysalis longicornis Neumann, 1901: a thesis presented in partial fulfilment of the requirements for the degree of Master of Veterinary Science at Massey University (Doctoral dissertation, Massey University). https://mro.massey.ac.nz/handle/10179/4944

Phillips, S. J., Anderson, R. P., & Schapire, R. E. (2006). Maximum entropy modeling of species geographic distributions. Ecological modelling, 190(3-4), 231–259. https://www.sciencedirect.com/science/article/pii/S030438000500267X

Pimentel, D., Lach, L., Zuniga, R. and D. Morrison. (2000) Environmental and Economic Costs of Nonindigenous Species in the United States. BioScience 50 (1), 53–65

R Core Team (2016). R: A language and environment for statistical computing. R Foundation for Statistical Computing, Vienna, Austria. URL https://www.R-project.org/.

Rainey, T., Occi, J. L., Robbins, R. G., & Egizi, A. (2018). Discovery of Haemaphysalis longicornis (Ixodida: Ixodidae) Parasitizing a Sheep in New Jersey, United States. Journal of Medical Entomology, 55(3), 757–759.

USDA-APHIS (2018) National Haemaphysalis longicornis (longhorned tick) Situation Report - August 29th, 2018.

USDA-NASS (2014). 2012 Census of Agriculture. https://www.agcensus.usda.gov/Publications/2012/

Vink, W. D., Lawrence, K., McFadden, A. M. J., & Bingham, P. (2016). An assessment of the herd-level impact of the Theileria orientalis (Ikeda) epidemic of cattle in New Zealand, 2012-2013: a mixed methods approach. New Zealand veterinary journal, 64(1), 48–54.

Watts, J. G., Playford, M. C., & Hickey, K. L. (2016). Theileria orientalis: a review. New Zealand veterinary journal, 64(1), 3–9. https://www.tandfonline.com/doi/full/10.1080/00480169.2015.1064792

Whitten, L. K. (1970). The control of external parasites of cattle. New Zealand veterinary journal, 18(7), 146–147.

Yu, Z. J., Lu, Y. L., Yang, X. L., Chen, J., Wang, H., Wang, D., & Liu, J. Z. (2014). Cold hardiness and biochemical response to low temperature of the unfed bush tick Haemaphysalis longicornis (Acari:). Parasites & vectors, 7(1), 346. https://parasitesandvectors.biomedcentral.com/articles/10.1186/1756-3305-7-346

Zheng, H., Yu, Z., Chen, Z., Zhou, L., Zheng, B., Ma, H., & Liu, J. (2011). Development and biological characteristics of Haemaphysalis longicornis (Acari: Ixodidae) under field conditions. Experimental and Applied Acarology, 53(4), 377–388. https://link.springer.com/article/10.1007/s10493-010-9415-3

